# Proximity to an SGC-DLPFC Individualized Functional Target and outcomes in large rTMS clinical trials for Treatment-Resistant Depression

**DOI:** 10.1101/2025.07.09.662866

**Authors:** Elizabeth Gregory, Shan H Siddiqi, Michael D Fox, Daniel M. Blumberger, Jonathan Downar, Zafiris J. Daskalakis, Katharine Dunlop, Fidel Vila-Rodriguez

**Affiliations:** Non-Invasive Neurostimulation Therapies (NINET) Laboratory, Department of Psychiatry, University of British Columbia, Vancouver, BC, Canada; Center for Brain Circuit Therapeutics, Brigham & Women’s Hospital, Boston, MA; Departments of Psychiatry, Pharmacology & Toxicology, and Institute of Medical Science, University of Toronto; Centre for Depression and Suicide Studies, St. Michael’s Hospital, Toronto, ON, Canada; Department of Psychiatry, School of Medicine, University of California, San Diego Health, San Diego, CA, United States; School of Biomedical Engineering, University of British Columbia, Vancouver, BC, Canada

**Author notes:** Correspondence to: Fidel Vila-Rodriguez, MD, PhD, FRCPC, DFAPA Non-Invasive Neurostimulation Therapies (NINET) Laboratory Department of Psychiatry, University of British Columbia 2255 Wesbrook Mall, Vancouver, BC V6T 2A1, Canada.

## Abstract

**Background:** Targeting methods for repetitive transcranial magnetic stimulation (rTMS) in patients with depression now include the use of individual functional scans to target specific functional connectivity (FC) patterns obtained from functional magnetic resonance imaging (fMRI). Potential biomarkers of rTMS response include target FC with the subgenual anterior cingulate cortex (SGC) or the causal depression circuit (CDC), each of which may be candidates for individualized functional targets (iFTs). We assessed the relationship of these two approaches to clinical outcomes in two large rTMS clinical trials.

**Methods:** 501 subjects with moderate to severe depression underwent 4-6 weeks of daily rTMS to the left dorsolateral prefrontal cortex (DLPFC), targeted using neuronavigation to a common group-based functional target. Resting-state scans acquired at baseline were used to retrospectively compute iFTs using either SGC-DLPFC or CDC-DLPFC FC. The Euclidean distance from the group-based target used in the trial to the centre of gravity of each iFT was computed and correlated with outcomes.

**Results:** Most subjects’ iFTs were within 2cm of their group-based target. Proximity to either the SGC- or CDC-iFT was not associated with better outcomes. Sensitivity analyses accounting for treatment target FC, methodology, data quality, or treatment parameters did not change the results.

**Conclusions:** Proximity to SGC- or CDC-derived iFTs was not associated with better outcomes in patients who received neuronavigated rTMS to a group-based target. Prospective randomized clinical trials comparing neuronavigated group-based target to neuronavigated iFTs are needed.

## INTRODUCTION

Repetitive transcranial magnetic stimulation (rTMS) is a first-line intervention for the management of treatment-resistant depression (TRD)(1), typically targeting the left dorsolateral prefrontal cortex (L-DLPFC)(2), achieving response and remission rates of 40-60%, and 20-40%, respectively(1).

rTMS induces an approximately 2-3cm-wide electrical field (e-field) within the cerebral cortex(3), and this local stimulation leads to modulation of brain circuits beyond the treatment target(4,5); thus, functional connectivity (FC) of the treatment target with distal brain regions may be particularly relevant in how rTMS exerts its antidepressant effects(4,6). FC between the DLFPC and the subgenual anterior cingulate cortex (SGC), a limbic region associated with hyperactivity in depression and believed to play a critical role the pathophysiology of the illness(7), is of particular interest in the context of rTMS(8). In group-level analyses, L-DLPFC treatment targets showing greater anticorrelation with the SGC are associated with more favorable outcomes, thereby suggesting that targeting this FC pattern could lead to increased response rates(9–11).

In 2012, Fox and colleagues proposed specific MNI coordinates as treatment targets, based on group-level analyses in healthy volunteers identifying regions within the L-DLPFC showing maximal SGC anticorrelation(9). This approach relies on the use of neuronavigation, where individual structural MRI head scans are registered in space with fiduciary markers, allowing for precise, consistent targeting of pre-specified coordinates(12). However, group-level targeting does not account for individual differences in FC. Instead, the use of an individualized functional target (iFT), which maximizes SGC-DLPFC anticorrelation at the individual level, may be key to achieving greater rates response to rTMS(10).

A number of studies using small, retrospective samples found that the distance between the clinically applied target and a retrospectively computed iFT, using individual fMRI data, was strongly associated with treatment response(13–16), thereby suggesting that delivering rTMS to an iFT could achieve significant gains in clinical outcomes. Prospectively, Cole and colleagues employed individualized targeting based on SGC anticorrelation paired with an accelerated protocol, reporting response rates of over 80%, notably higher than that reported by standard protocols. However, their study did not directly compare individualized targeting to other approaches and involved accelerated protocol; thus, the specific impact of delivering stimulation to iFTs is unclear(17).

In contrast, non-replication of the relationship between stimulation site anticorrelation and treatment outcomes suggests alternative patterns of FC may be superior targets for developing iFTs(18). The causal depression circuit (CDC), recently defined by Siddiqi et al., is a distributed map derived from both brain lesions and neurostimulation sites associated with depressive symptoms, with peaks in the DLPFC, SGC, and several other brain regions. The CDC demonstrates specificity to depression, and group-level connectivity of rTMS treatment sites with the CDC better predicted outcomes compared to connectivity with the SGC(19). Individualized targets identified using distributed circuit-maps are more reliable than those identified using a small SGC region (10) and thus individualized targets identified using the CDC may be a promising alternative(19,20).

The current study sought to examine the relationship between the proximity to an iFT and treatment response in a large, retrospective dataset of depressed patients that received rTMS. We computed iFTs using two approaches: 1) SGC-DLPFC FC (SGC-iFT) and 2) CDC-DLPFC FC (CDC-iFT), using previously validated computational methods(13,20,21). We sought to replicate previous findings that proximity to a SGC-seeded iFT would predict treatment outcomes in a large sample from two clinical trials using neuronavigation. In addition, we investigated whether a CDC-derived iFT would similarly predict treatment outcomes.

## METHODS

### Dataset

Analyses were conducted in two cohorts of patients with major depressive disorder (MDD) recruited to two rTMS clinical trials across three Canadian sites(2,22). All participants received rTMS to the L-DLPFC for 4-6 weeks at 120% resting motor threshold (RMT) intensity. Treatment was delivered to MNI coordinates [-38, 44, 26], corresponding to an optimized SGC group-target(9), using the *Visor2* neuronavigation system (ANT Neuro, Enschede, Netherlands). Prior to rTMS, all subjects underwent a T1-weighted scan and a 10-minute single-echo resting-state fMRI (rs-fMRI) scan (300 volumes). Further details on cohorts, including scanning parameters, are provided in the supplement (section 1.1).

### fMRI preprocessing

fMRI data was preprocessed using fMRIPrep (version 23.2.0)(23); details are presented in the supplement (section 1.2). The fMRIPrep output then underwent removal of non-steady state volumes, spatial smoothing with a 6mm FWHM gaussian kernel, and non-aggressive motion artifact extraction using ICA-AROMA(23). The white matter, CSF, and global signal time-series generated by the fMRIPrep workflow were regressed from the data, followed by bandpass filtering.

### Time Series Derivation

Individual timeseries were calculated for each voxel within the L-DLPFC, which was defined by combining 20mm radius spheres centered at BA9 (MNI –36, 39, 43), BA46 (MNI –44, 40, 29), the average ‘5cm’ TMS site (MNI –41, 16, 54), and the average BeamF3 site (MNI –37, 26, 49)(21). The SGC timeseries was derived based on the previously defined seedmap approach(21), which leverages signal from all grey matter voxels to estimate SGC connectivity, using each voxel’s weight in a group-level seedmap of the right SGC. Timeseries for the CDC was created using a similar method, weighting the signal from each gray matter voxel by the combined CDC *r* map(19). Voxels within the L-DLPFC ROI were excluded when generating both timeseries. Further details are described in the supplement (section 1.3).

### iFT Localization

While multiple approaches exist for localizing iFTs (17,24,25), the cluster-based approach has been reported to minimize intraindividual variation (*i.e.* reliability within subjects), while maximizing interindividual spatial variation, for iFTs derived from SGC-DLPFC FC (21) and is similar to the approach used by Cole et al in the accelerated iTBS trials (17). We delineated clusters (defined using standard 26-voxel neighborhoods) within the top 0.5% of most anticorrelated voxels, per previous work identifying this threshold as optimal for seedmap-generated data(21), then selected the centre of gravity of the largest cluster as the iFT. We also computed targets using higher cluster thresholds (1%, 2.5%, 5%, and 10%) as well as an alternative “searchlight method (supplement, section 2.1).

The Euclidean distance between the iFT and applied group-target was calculated in subject space.

### Outcome Measures

Clinical outcome was operationalized as the percent change in HRSD-17 scores from baseline to end of treatment, which was the primary outcome reported across cohorts. We also assessed response (HRSD-17≥50% reduction) and remission (HRSD-17<8). (26)

The relationship between the Euclidean distance to the iFT (SGC or CDC) and clinical outcomes was assessed using correlation and independent samples t-tests. Target FC was also computed using a 12mm weighted cone (9)to adjust for any effects of stimulation site connectivity on outcomes. Using the Shapiro-Wilk test, measures for distance, target FC, and percent change in HRSD-17 scores violated assumptions of normality (W=0.83-0.97, *p*<.001 for all). Thus, unless stated otherwise, Spearman’s rho was used in all correlation analyses.

### Comparison of targeting approaches

We assessed the similarity of the approaches used to derive iFTs through spatial correlations of the original SGC seedmap/CDC *r* maps used to derive timeseries, spatial correlations of subject-level FC maps of the SGC and CDC timeseries, as well as computing intraindividual distances between iFTs.

(13–15)**Error! Bookmark not defined.**

## RESULTS

525 subjects completed treatment and had baseline scans. 24 subjects were excluded due to high framewise displacement (supplement, section 1.2.4), leaving 501 subjects for analysis. Sample characteristics are summarized in Table 1.

**Table 1.** Summary of clinical and demographic variables of the analytic sample (N=501), stratified by cohort.

|  | **THREE-D** | **CARTBIND** |
| --- | --- | --- |
|  | n = 337 | n = 164 |
| **Measure** | **Mean (SD) or count (%)** | **Mean (SD) or count (%)** |
| Age ; years | 42.45 (11.44) | 41.43 (11.30) |
| Number of years of education | 16.40 (2.99) | 17.91 (2.32) |
| Sex; Female | 200 (59%) | 107 (65%) |
| Handedness; right | 302 (89%) | 147 (90%) |
| Baseline HRSD<sup>a</sup> score | 23.27 (4.28) | 23.18 (3.89) |
| End of rTMS HRSD score | 12.82 (7.72) | 13.49 (6.59) |
| % Change in HRSD score | -44.82 (32.24) | -41.06 (27.82) |
| Responder<sup>b</sup> | 171 (50%) | 67 (41%) |
| Remitter<sup>c</sup> | 111 (33%) | 35 (21%) |
| Age of onset; years | 20.82 (10.87) | 20.67 (9.46) |
| Length of current episode; months | 23.07 (27.68) | 36.66 (50.56) |
| Anxiety comorbidity<sup>d</sup> | 175 (51%) | 89 (54%) |
| ATHF<sup>e</sup> score | 6.29 (3.35) | 8.05 (5.23) |
| Benzodiazepines | 110 (32%) | 41 (25%) |
| Antidepressants | 263 (77%) | 99 (60%) |
| Antipsychotics | 57 (17%) | 35 (21%) |
| Received Theta-burst stimulation | 177 (52%) | 164 (100%) |
| rTMS stimulation intensity (% MSO) | 49.77 (10.88) | 50.85 (10.98) |
| Number of daily treatments | 26.72 (4.68) | 29.99 (0.11) |

<sup>a</sup>Hamilton Rating Scale for Depression, 17 item version.
<sup>b</sup>Response defined as a 50% or greater decrease in HRSD score at the end of treatment.
<sup>c</sup>Remission defined as an HRSD score of 8 or less at the end of treatment.
<sup>d</sup>Per Mini International Neuropsychiatric Interview performed at screening visits.
<sup>e</sup>Antidepressant treatment history form.

### Individualized targeting

The median distance from the treatment target to the SGC-iFT was 12.8mm (SD=12.9), and to the CDC-iFT, 13.8mm (SD=13.4). A majority of iFTs fell within 20 mm of the treatment target (SGC-iFT: 74%; CDC-iFT: 69%); there was no difference in HRSD change for subjects with iFTs within, versus beyond, 20 mm (SGC-iFT: t(209.60)=0.64, *p*=0.5; CDC-iFT: t(286.09)=1.23, *p*=0.2).

HRSD change was not significantly correlated with SGC-iFT distance (ρ=0.02, *p*=0.6), nor CDC-iFT distance (ρ=0.05, *p*=0.3). There was also no significant difference in distance to either iFT between responders and non-responders (SGC-iFT: *t*(497.34)= -0.11, *p*=0.9; CDC-iFT *t*(498.99)= -1.16, *p*=0.2), or remitters and non-remitters (SGC-iFT: *t*(273.07)=0.15, *p=*0.9; CDC-iFT: *t*(286.56)= -1.86, *p*=0.06).

FC at the group-target was also not correlated with HRSD change for both the SGC (ρ = 0.04, *p*=0.4) and the CDC (ρ= -0.03, *p*=0.5).

Sensitivity analyses revealed no effect of proximity on outcomes when computing iFTs at a range of cluster thresholds or using the “searchlight method” (supplement, 2.1). Additionally, we found no effect of signal quality (either whole-brain or within the SGC), scanner site, cohort, or treatment assignment (supplement, 2.2-2.5). Notably, there were differences in the overall metrics of proximity and target FC across scanner sites and cohorts, suggesting iFT computation was affected by scanner and/or scanning protocol (supplement, 2.3).

### Comparison of SGC and CDC individualized targeting approaches

The SGC seedmap and CDC R map (used to derive timeseries) themselves showed a moderate negative spatial correlation of *r*=-0.54 (*p*<.001). The timeseries derived from these maps showed a strong within-subject negative correlation (M*=*-0.85, SD=0.18).

Whole-brain FC maps of the SGC and CDC were computed and compared within-subject (Figure 3a and 3b). At the subject level, SGC and CDC whole-brain FC maps showed a strong negative correlation, with a mean *r* value of -0.84 (SD=0.29). Within the L-DLPFC, FC maps showed a mean spatial correlation of -0.85 (SD=0.29). The calculated FC at the treatment target itself also showed a strong correlation between the SGC and CDC approaches (ρ*=* -0.89*, p<.*001).

**Figure 1.**
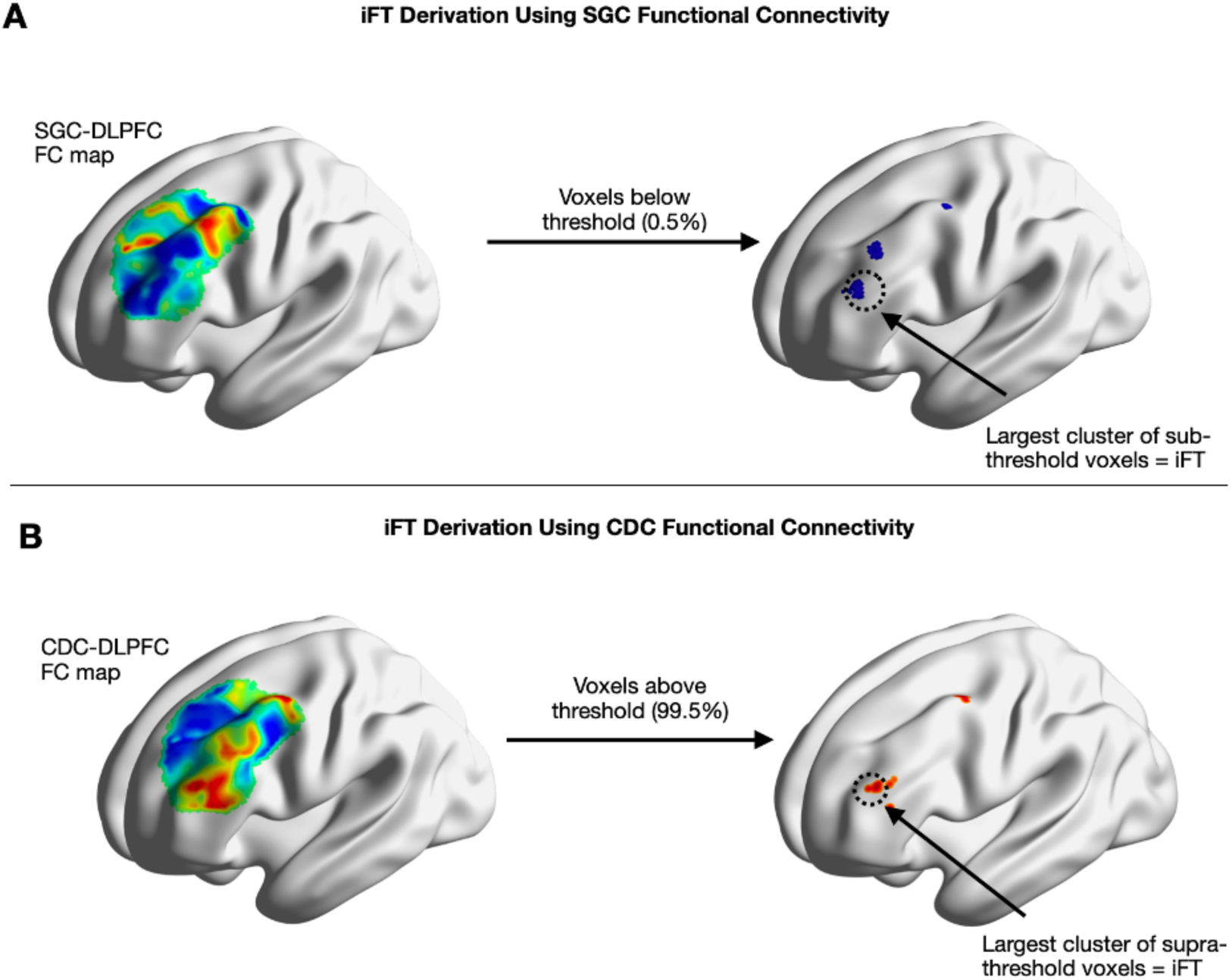
Methods for computing the location of the iFT for each subject, as demonstrated in a representative subject. A) SGC-DLPFC functional connectivity was computed by taking the SGC seedmap timeseries and correlating it with each timeseries in the DLPFC mask. Next, the 0.5% most anti-correlated voxels (below 0.5%) were identified, and the largest cluster of sub-threshold voxels was selected as the optimal cluster. B) CDC-DLPFC functional connectivity was computed by taking the CDC seedmap timeseries and correlating it with each timeseries in the DLPFC mask. Next, the 0.5% most correlated voxels (above 99.5%) were identified, and the largest cluster of supra-threshold voxels was selected as the optimal cluster. In both scenarios, iFT was computed by taking the centre of gravity of the largest cluster.

**Figure 2.**
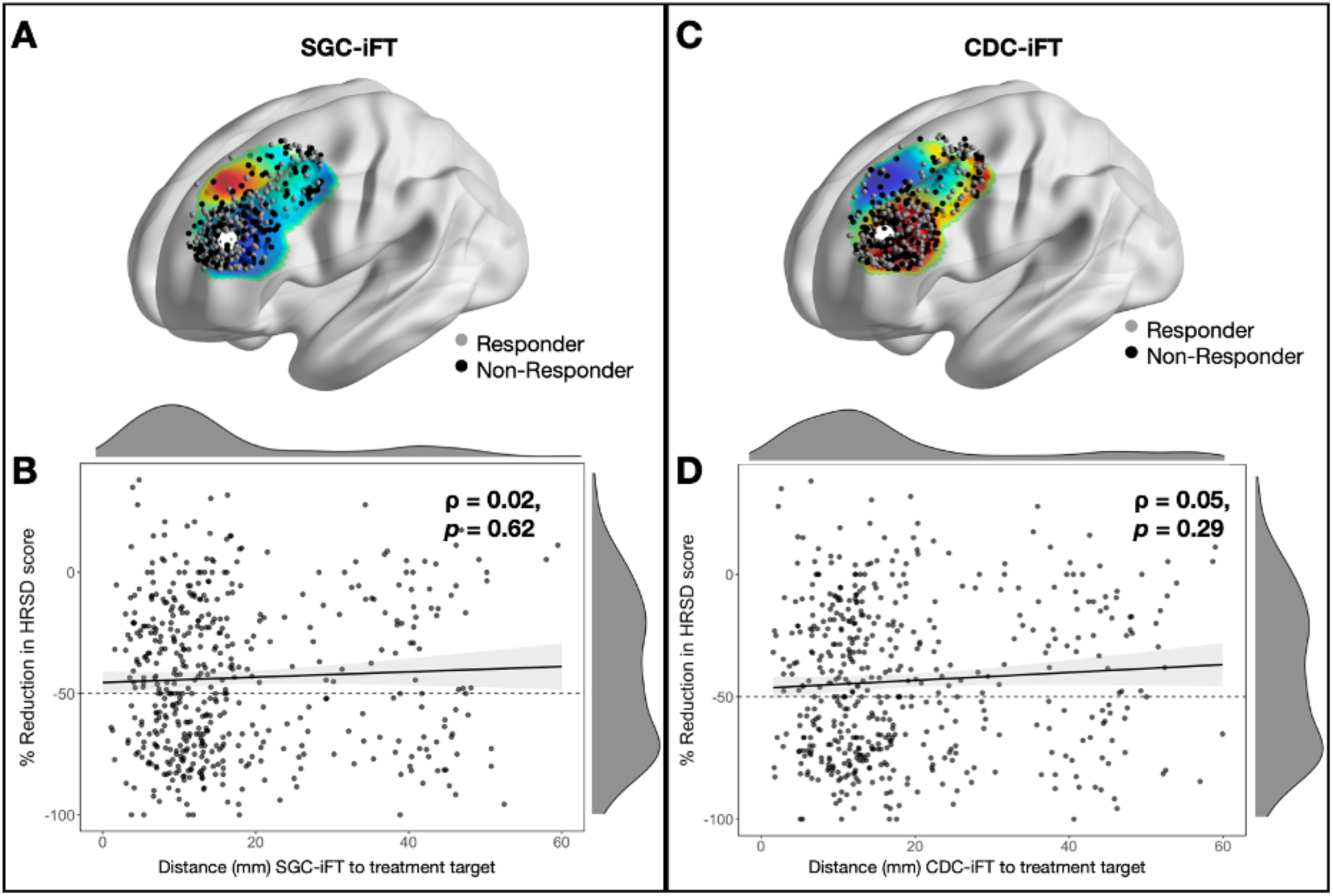
Topographical distribution of optimal targets and their proximity to the actual treatment target. For the SGC-iFT approach, (A) shows the distribution of individualized targets within the DLPFC (mapped over mean FC with the SGC seedmap timeseries) and (B) plots the relationship between iFT proximity and treatment response. Similarly, for the CDC-iFT approach, (C) shows the topographical distribution of individualized targets (mapped over the mean FC with the CDC timeseries) while (D) plots the relationship between iFT proximity and treatment response. For (A) and (B), grey circles represent responders, black circles represent non-responders, and the treatment target is denoted by the larger white circle. **ρ** (rho), ***p*** (p value)

**Figure 3.**
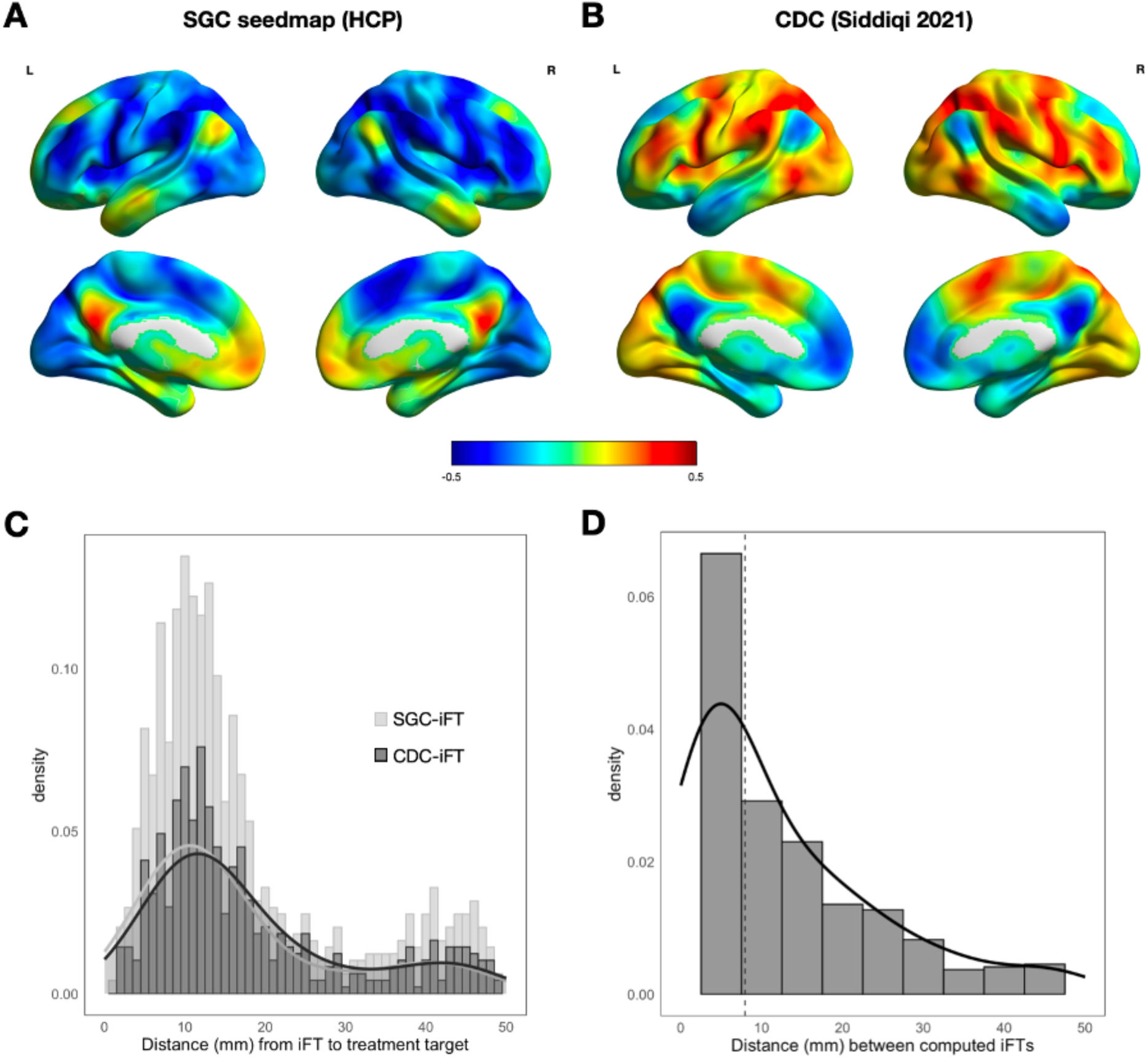
Comparison between the iFT derivation methods. Mean whole-brain FC maps were computed for both the SGC-iFT (A) and CDC-iFT approaches (B), which show a strong inverse spatial correlation (M=-0.84, SD = 0.28). (C) The distribution of distances derived from each iFT approach are shown as stacked histograms. (D) Distribution of the distance between each subject’s SGC-iFT and CDC-iFT. Vertical dashed line represents the median (equal to 7.94).

Subjects’ SGC and CDC-derived iFTs were close to each other for most participants, with a median distance of 7.9mm (SD=13.7) (Figure 3c). In terms of the distance measure itself, iFT distance distributions were statistically different (t(500)=3.03, *p=*0.002, 95% CI=0.56 - 2.62) between the two methods (Figure 3d), though the distances measures from the two iFTs were moderately correlated (ρ=0.57, *p*<0.001).

## DISCUSSION

In this large sample of patients with TRD receiving rTMS to a neuronavigated group-based target, we did not find an association between proximity to an SGC-iFT and treatment outcome. These results contrast with the positive results in three smaller-scale studies, but converge with recent work by Elbau and colleagues reporting that SGC FC contributes only marginally to treatment outcomes within a subsample of the THREE-D dataset(13–15,20). Our comprehensive analyses encompassed not only these subjects, but also extended to a subsample of THREE-D scanned at a different site, as well as to the multi-site CARTBIND dataset.

We additionally calculated iFTs using the CDC, finding similarly negative results. Siddiqi and colleagues previously found that treatment site-CDC FC explained more variance in response compared to SGC FC(19). As a circuit-based approach, deriving iFTs using the CDC could be, in theory, a more robust targeting method(27). However, given the SGC acts as a node of the CDC, and given patients in this analysis received stimulation to a target based on group-level SGC FC, this may have decreased the variation needed to capture any effect.

### Use of neuronavigation and group-based Targeting

In our sample, the targeted coordinates were previously identified as showing maximal group-level SGC anticorrelation(10). All subjects in the current analysis received rTMS to this target with neuronavigation, which achieves a fair level of targeting accuracy, typically within 4-8mm(12), as well as enhanced consistency in coil placement and orientation relative to scalp-based heuristic methods(28). One interpretation is that individual differences in SGC anticorrelation are *most likely* to be observed near this target and the e-field is thus likely to include the iFT. The e-field from a butterfly coil, per real-head finite element boundary simulations, is estimated to span several cm^2^(3). Median distances for SGC- and CDC-derived iFTs were approximately 12-13mm, with most iFTs within 20mm of the stimulated group-target, which is similar to a prior study which employed this same group-target {Citation}. In contrast, iFTs were further from average MNI coordinates of other commonly used targets (5cm and BeamF3), with these alternate targets also showing weaker FC with SGC and CDC timeseries (supplement, 2.7). Because the stimulated target was close to most subjects’ iFTs, it’s possible that we lacked the statistical variation needed to detect an effect of iFT proximity.

Our results are consistent with the recent BRIGhTMIND trial where no differences were found between a neuronavigated group-based target and an iFT (albeit seeded from insula rather than SGC). To our knowledge this is the only head-to-head comparison of neuronavigation to a group-target versus iFT published to-date, though interpretation and translation of those findings to our data are complicated by treatment group differences beyond the targeting approach (*e.g*. stimulation protocol)(29).

### Potential effects of treatment parameters beyond targeting

Targeting is only one aspect of rTMS individualization. The vast parameter space in rTMS poses a challenge in disentangling the relative contributions of any one parameter of treatment delivery, as well as interpreting differences in findings between studies.

Compared to the previous small studies finding positive results, one key difference is that a majority of our sample received iTBS stimulation, while HFL protocols were exclusively delivered in the previous publications(13–15). However, analyses revealed similarly null results within iTBS- and HFL-treated groups (supplement, Figure S6), thus suggesting findings cannot be explained by differences in the frequency parameter.

Stimulation intensity, delivered at 120% rMT in both our cohorts, is an additional treatment parameter to consider. While standard, and similar to doses reported in previous studies finding a relationship between iFT proximity and outcomes (110-120% rMT; (14–16,19), this dose is considerably higher than conventional iTBS dosing to the motor cortex, typically 70-80% rMT or active motor threshold(30,31). In the SNT trial, Cole and colleagues employed a 90% rMT scalp-to-cortex distance corrected dose (achieving a mean dose of 102% rMT), reporting excellent response rates to accelerated iTBS(17). (14,15,19)While the question of optimal stimulation dose is beyond the scope of this current work, it is worth considering that suprathreshold stimulation means that a larger volume of the cortex is stimulated at subthreshold levels, leading to neurobiological effects upon cortical areas more distal to the stimulation target, effectively reducing focality(32).

Electrical field (e-field) variability is another potential consideration. Target placement on a gyral crown versus a sulcal groove, as well as the coil orientation relative to the cortical folding, affect whether the selected cortical region is sufficiently stimulated(33); thus, targeting a set of pre-defined coordinates can still lead to significant between-subject differences(34).

Individual differences in e-field may have obfuscated our ability to detect significant effects of iFT proximity. However, previous publications reporting a relationship between iFT proximity and outcomes similarly employed standard targeting without e-field optimization(13–15). A final consideration with respect to the e-field is the choice of TMS coil, due to variations in focality and depth(35); all subjects in the current analysis received stimulation with a lesser focal coil (MagVenture B70).

Lastly, preclinical studies have reported ‘non-response’ even when stimulation is known to be on-target: for example, in M1, more than half of participants showed minimal facilitation of motor evoked potentials despite multiple sessions of iTBS(31). Such observations suggest that insufficient plasticity, rather than off-target stimulation, may underlie non-response for many patients. In keeping with this possibility, recent work suggests that enhancing plasticity with D-cycloserine can markedly increase response and remission rates, even with relatively short TMS courses of 20 sessions without individualized targeting(36).

### Considerations of heterogeneity in depression

MDD is a highly heterogeneous disorder, with diverse underlying FC patterns contributing to morbidity across patients. Targeting SGC FC may offer therapeutic benefits to a specific subset of MDD patients, while for others, alternate patterns of FC may serve as more promising candidates. Dunlop *et al.* delineated four distinct biotypes, characterized by variations in symptomatology, FC profiles, and response to rTMS(37). Further, discrete clusters of symptoms, namely dysphoric and anxiosomatic, have been found to respond to rTMS through two distinct circuits(38). Thus, targeting SGC FC may offer therapeutic benefits to a specific subset of MDD patients, while for others, alternate patterns of FC may serve as more promising candidates. Post-hoc analysis of symptom cluster change (mood, anxiety, somatic, sleep), however, found no significant association with measures of proximity or target FC (supplement, 2.6).

### Limitations: considerations of fMRI scan quality

Resting-state fMRI images were acquired on 3T scanners, with single-echo imaging, larger-than-typical voxel sizes, and no fieldmap acquisition. These now legacy scan parameters are critical to consider when appraising our findings.

Previous investigation of the methods employed in the current analysis (*ie* cluster-based iFT localization, seedmap approach for SGC FC) found that optimal target localization was achieved with at least 20minutes of scan time in the HCP dataset(21). Other work estimates a minimal scan duration for single-echo within the range of 10-40 minutes to produce reliable iFTs within the DLPFC(39). Given the relatively short scanning duration of 10minutes (TR=2s, 300 volumes), derived iFTs may have been unstable. Prior studies reporting positive results were all conducted with single-echo acquisition protocols, with varying amounts of resting-state data: 6 minutes (TR= 2.05s, 176volumes)(16), 6.5minutes (TR=2.5s, 149 volumes)(15), 13.33minutes (TR=2s, 400volumes)(14), and 28minutes (TR=3.2s, 525volumes)(13).

The acquired voxel dimensions varied by cohort but were larger than typical dimensions in all cases (THREE-D UHN & CAMH: 3.44×3.44×5.0mm; THREE-D UBC: 2.75×2.75×5.0; CARTBIND, all sites: 4.0×4.0×4.0mm). Larger voxels may benefit from increased tSNR, albeit with the trade-off of reduced spatial precision. Previous positive findings were reported from studies with acquired voxel dimensions of 2.1×2.1×2.8mm(16), 3.5×3.5×3.5mm(14,15), or 3.75×3.75×3mm(13). Thus, an outstanding question is whether larger voxels in our cohorts can explain the null findings. We found that iFT proximity was influenced by both cohort and scanner site (supplement, section 2.3), suggesting an effect not only of acquisition parameters but also scanner-specific differences, given identical acquisition parameters across sites for the CARTBIND trial. However, we did not find any relationship between iFT proximity and outcomes within any scanner or scan protocol. This factor should be considered when interpreting our findings, and in any prospective application of iFTs, as iFT computation may be sensitive to scan parameters.

A critical consideration in the fMRI imaging sequences used is the impact on signal within the SGC, which is associated with significant signal drop-out in single-echo scans due to proximity to the sinuses(40). To attempt to mitigate this issue in our dataset, we employed a seedmap approach, which is a well-validated method to reliably estimate SGC FC(10,20). We probed the effect of SGC signal dropout in our sample by computing tSNR, finding no significant effects within subjects with higher whole-brain or SGC tSNR (supplement, 2.2). Future study on this topic, including using data collected with alternative scan techniques (such as multi-echo), longer acquisition times, and smaller voxels, would be of benefit to understand the reasons for the null findings in this case.

Lastly, multiple approaches exist for computing iFTs, including targeting alternate patterns of FC (29) or employing task-fMRI to achieve more reliable cortical activations (41). Though beyond the scope of this work, it is possible that alternate approaches may be more efficacious in personalizing rTMS targeting for depression.

### Conclusion

This retrospective analysis of 501 subjects receiving neuronavigated high-frequency rTMS to the L-DPFC did not find any effect of proximity to an individualized target on treatment outcomes to. These findings should be considered in the context of the quality of the fMRI scans and specific methods used to derive iFTs. Ultimately, a prospective randomized clinical trial may be necessary to determine the clinical value of such targeting approach.

## Acknowledgements

We are grateful for Dr. Robin Cash and Dr. Andrew Zalesky for sharing code to derive individualized functional targets and invaluable discussion on the manuscript.

## Author contributions

Conceptualization: EG, SHS, FVR; Formal analysis: EG; Methodology: EG, SHS, MDF, KD, FVR; Investigation: FVR, DM, JD, ZJD; Funding Acquisition: DM, JD, FVR, ZJD; Writing – Original Draft: EG, FVR; Writing – Review & Editing: EG, SHS, MDF, DMB, JD, ZJD, KD, FVR; Supervision: FVR

## Funding

This work was supported in part by Brain Canada with financial assistance from Brain Canada, Canadian Institutes of Health Research MOP-136801, Vancouver Coastal Health Research Institute, the Michael Smith Foundation for Health Research, as well as Philanthropic sources. MDF was supported by grants from the NIH (R01MH113929, R21MH126271, R21NS123813, R01NS127892, R01MH130666, UM1NS132358), the Kaye Family Research Endowment, the Ellison / Baszucki Family Foundation, the Once Upon a Time Foundation, the Manley Family, Donna and Tom May, and Boston Scientific.

## Disclosures

DMB receives research support from CIHR, NIMH, Wellcome Trust, Brain Canada and the Temerty Family through the CAMH Foundation and the Campbell Family Research Institute, has received research grant support and in-kind equipment support for an investigator-initiated study from Brainsway Ltd, was the site principal investigator for three sponsor-initiated studies for Brainsway Ltd, has received in-kind equipment support from Magventure for investigator-initiated studies, has received medication supplies for an investigator-initiated trial from Indivior, is a scientific advisor for Sooma Medical and is the Co-Chair of the Clinical Standards Committee of the Clinical TMS Society (unpaid). EG holds a Vanier Canada Graduate Scholarship from the Canadian Institutes of Health Research (CIHR), as well as stipend support from the Brain Canada Foundation. FVR has received research support from CIHR, Brain Canada, Michael Smith Foundation for Health Research, Vancouver Coastal Health Research Institute, and Weston Brain Institute for investigator-initiated research. Philanthropic support from Seedlings Foundation. In-kind equipment support for investigator-initiated trial from MagVenture. He has received honoraria for participation in an advisory board for Allergan. FVR is a volunteer director on the board of directors of the British Columbia Schizophrenia Society.

He is a member of the Education Committee of the Clinical TMS Society (unpaid). JD has received research support from NIH, CIHR, Brain Canada, Ontario Brain Institute, the Klarman Family Foundation, the Arrell Family Foundation, and the Buchan Family Foundation, in-kind equipment support for investigator-initiated trials from MagVenture, is an advisor for BrainCheck, Arc Health Partners and Salience Neuro Health, and is a co-founder of Ampa Health. KD holds an Academic Scholars Award from the Department of Psychiatry, University of Toronto, and is listed as an inventor for Cornell University patent applications on neuroimaging biomarkers for depression that are pending or in preparation. KD has also received research support from the American Foundation for Suicide Prevention, the Brain & Behavior Research Fund, Brain Canada, CIHR, and the St. Michael’s Foundation.

MDF has intellectual property on the use of brain connectivity imaging to analyze lesions and guide brain stimulation, has consulted for Magnus Medical, Soterix, Abbott, Boston Scientific, Tal Medical, and MDC Venture Capital, has received research support from Neuronetics and Boston Scientific, and sits of the SAB of Salma Health. SHS serves as a consultant for Kaizen Brain Center, Acacia Mental Health, and Magnus Medical. SHS owns stock in Brainsway Ltd (publicly traded) and Magnus Medical (not publicly traded). SHS has received investigator-initiated research support from Neuronetics Inc and Brainsway Ltd.

