## Supplement for "Proximity to an SGC-DLPFC Individualized Functional Target and outcomes in large rTMS clinical trials for Treatment-Resistant Depression"

### SUPPLEMENTAL INFORMATION

### 1. METHODS

#### 1.1 Cohorts

Both cohorts are clinical trials that were conducted at three Canadian academic health centres: Centre for Addiction and Mental Health, Toronto, ON; University Health Network, Toronto, ON; University of British Columbia Hospital, Vancouver, BC).

The inclusion and exclusion were identical for both clinical trials, with the exception of age, which was between 18-65 for THREE-D and 18-59 for CARTBIND, and the degree of treatment resistance (in THREE-D, less than four adequate antidepressant trials). The additional criteria were as follows:

**Inclusion Criteria:**

1. are outpatients
2. are voluntary and competent to consent to treatment
3. have a Mini-International Neuropsychiatric Interview (MINI) confirmed diagnosis of MDD, single or recurrent
4. have failed to achieve a clinical response to an adequate dose of an antidepressant based on an Antidepressant Treatment History Form (ATHF) score for that antidepressant trial of > 3 in the current episode OR have been unable to tolerate at least 2 separate trials of antidepressants of inadequate dose and duration (ATHF score of 1 or 2 on those 2 separate antidepressants)
5. have a score >= 18 on the HRSD-17 item
6. have had no increase or initiation of any psychotropic medication in the 4 weeks prior to screening
7. able to adhere to the treatment schedule
8. Pass the TMS adult safety screening (TASS) questionnaire
9. have normal thyroid functioning based on pre-study blood work.

**Exclusion Criteria:**

1. have a Mini-International Neuropsychiatric Interview (MINI) confirmed diagnosis of substance dependence or abuse within the last 3 months
2. have a concomitant major unstable medical illness, cardiac pacemaker or implanted medication pump
3. have active suicidal intent
4. are pregnant
5. have a lifetime Mini-International Neuropsychiatric Interview (MINI) diagnosis of bipolar I or II disorder, schizophrenia, schizoaffective disorder, schizophreniform disorder, delusional disorder, or current psychotic symptoms
6. have a MINI diagnosis of obsessive-compulsive disorder, post-traumatic stress disorder (current or within the last year), anxiety disorder (generalized anxiety disorder, social anxiety disorder, panic disorder), or dysthymia, assessed by a study investigator to be primary and causing greater impairment than MDD
7. have a diagnosis of any personality disorder, and assessed by a study investigator to be primary and causing greater impairment than MDD
8. have failed a course of ECT in the current episode or previous episode
9. have received rTMS for any previous indication due to the potential compromise of subject blinding

10. have any significant neurological disorder or insult including, but not limited to: any condition likely to be associated with increased intracranial pressure, space occupying brain lesion, any history of seizure except those therapeutically induced by ECT or a febrile seizure of infancy, cerebral aneurysm, Parkinson's disease, Huntington's chorea, multiple sclerosis, significant head trauma with loss of consciousness for greater than 5 minutes
11. have an intracranial implant (e.g., aneurysm clips, shunts, stimulators, cochlear implants, or electrodes) or any other metal object within or near the head, excluding the mouth, that cannot be safely removed
12. if participating in psychotherapy, must have been in stable treatment for at least 3 months prior to entry into the study, with no anticipation of change in the frequency of therapeutic sessions, or the therapeutic focus over the duration of the study
13. clinically significant laboratory abnormality, in the opinion of the one of the principal investigators or study physicians

##### 1.1.1 THREE-D Cohort

The THREE-D clinical trial (clinicaltrials.gov ID NCT01887782) established non-inferiority of intermittent theta-burst (iTBS) to 10Hz repetitive transcranial magnetic stimulation (rTMS) to the left dorsolateral prefrontal cortex (L-DLPFC)(1). The trial was conducted from September 2013 to October 2016. In addition to clinical trial participants, an additional 20 subjects were included within the THREE-D cohort, drawn from a pilot trial conducted at the UBC site with identical parameters to the clinical trial (clinicaltrials.gov ID NCT02800226).
rTMS was delivered at 120% rMT, using either the MagPro X100 or R30 stimulator equipped with a B70 fluid-cooled coil and high-performance cooler (MagVenture, Farum, Denmark). RMT was determined prior to the first treatment according to standard clinical practice (2). The stimulation target was specified by reverse co-registration from a stereotaxic coordinate on the standard Montreal Neurological Institute (MNI-152) template brain on to each subject's anatomical (T1) MRI. Neuronavigation was performed using the *Visor 2* system (Advanced Neuro Therapeutics, Enschede, Netherlands) for each treatment session, positioning the coil for maximal field strength at MNI coordinates [-38, 44, 26].
10Hz rTMS was delivered 4 seconds on and 26 seconds off, 3000 pulses per session, and a total duration of 37.5 minutes. iTBS rTMS was delivered using triplet 50Hz bursts repeated at 5Hz, 2 seconds on and 8 seconds off, 600 pulses per session, and a total duration of 3 minutes 9 seconds. Participants received once-daily sessions on weekdays (5 sessions per week) for 20 sessions in total (4 weeks). After 20 sessions, participants whose HRSD-17 scores improved by more than 30% from baseline but who had not achieved remission, received an additional 10 once-daily sessions for a total of 30 sessions (6 weeks).
The scanning parameters for the THREE-D cohort were as follows:

1. **UHN**: Subjects were scanned on a GE Signa 3.0T scanner. The T1-weighted scan used a fast spoiled gradient echo (SPGR) sequence (TI = 300 ms, TE = minimum full, flip angle 20 degrees, 1.05x0.94x0.94mm voxels, FOV=240, 120 sagittal slices with interleaved acquisition). The resting-state fMRI scan used a gradient echo pulse sequence (TR = 2000 ms, TE = 30 ms, flip angle = 85 degrees, 3.44x 3.44 x5.0 mm voxels, FOV 220 x 220 x 160, 32 axial slices).
2. **UBC:** Subjects were scanned on a Philips Achieva 3.0T scanner. The T1-weighted scan used a FFE multi-shot single echo (TR = 8.1 ms, TE = minimum, flip angle 8 degrees, 1.00 x 1.02 x 1.00 mm voxels, FOV = 256x256x165, 165 sagittal slices with interleaved acquisition). The resting-state fMRI scan used a FFE single-shot EPI sequence (TR =

2000 ms, TE = 30 ms, flip angle = 90 degrees, voxel size = 2.75 x 2.75 x 5.0 mm voxels with 1 mm slice gap, FOV = 220 x 220 x 155, 26 axial slices with interleaved acquisition).

386 subjects received baseline fMRI scans. 353 subjects completed treatment and post-treatment clinical ratings. 16 subjects had excessive FWD (>0.3mm average), leaving 337 subjects remaining for analysis: 289 collected at UHN, 48 collected at UBC.

**1.1.2 CARTBIND Cohort**

The CARTBIND clinical trial (clinicaltrials.gov ID NCT02729792) randomized TRD to once or twice daily iTBS to the L-DLPFC for 6 weeks, finding no difference in rates of improvement(3). The trial was conducted from April 2016 to February 2018.

rTMS was delivered at 120% rMT, using either the MagPro X100 or R30 stimulator equipped with a B70 fluid-cooled coil and high-performance cooler (MagVenture, Farum, Denmark). RMT was determined prior to the first treatment according to standard clinical practice[23]. The stimulation target was specified by reverse co-registration from a stereotaxic coordinate on the standard Montreal Neurological Institute (MNI-152) template brain on to each subject's anatomical (T1) MRI. Neuronavigation was performed using the *Visor 2* system (Advanced Neuro Therapeutics, Enschede, Netherlands) for each treatment session, positioning the coil for maximal field strength at MNI coordinates [-38, 44, 26].

iTBS was delivered using the same parameters as THREE-D, though participants received 1200 pulses in total per day (as opposed to 600 pulses per day). Participants in the once daily group received 1200 pulses at a single session on weekdays for a total of 30 sessions over 6 weeks. Participants in the twice daily group received 1200 pulses of iTBS over two sessions per day, approximately 54 minutes apart (600 pulses per session), for a total of 60 sessions over 6 weeks.

203 subjects received baseline fMRI scans. 172 subjects completed treatment and post-treatment clinical ratings. 8 subjects had excessive FWD (>0.3mm average), leaving a total of 164 subjects for analysis – 40, 44, and 80 from UBC, UHN, and CAMH, respectively.

The scanning parameters for the CARTBIND cohort were as follows:

1. **UHN:** Subjects were scanned on a GE Signa 3.0T scanner. The T1-weighted scan used a BRAVO sequence (TR = 7.5 ms, TE = 2.9/2 ms, flip angle = 15 degrees, voxel size = 1.0x1.0x1.0mm, FOV = 240 mm, 180 sagittal slices with interleaved acquisition). The resting-state fMRI scan used a gradient echo sequence (TR = 2000ms, TE = 30 ms, flip angle = 75 degrees, voxel size = 4.0 x 4.0 x 4.0 mm, FOV = 256mm, 36 axial slices with interleaved acquisition).
2. **UBC:** Subjects were scanned on a Philips Achieva 3.0T scanner. The T1-weighted scan used a FFE sequence (TR = 6.5 ms, TE = 3 ms, flip angle 8 degrees, voxel size = 1.0x1.0x1.0mm, FOV=240 mm, 180 sagittal slices with interleaved acquisition). The resting-state fMRI scan used a FFE sequence (TR = 2000 ms, TE = 30 ms, flip angle = 75 degrees, voxel size = 4.0 x 4.0 x 4.0 mm, FOV=256 mm, 36 axial slices with interleaved acquisition).
3. **CAMH:** Subjects were scanned on a GE Discovery 3.0T scanner. The T1-weighted scan used a BRAVO sequence (TR = 7.5 ms, TE = 2.9/2 ms, flip angle = 15 degrees, voxel size = 1.0x1.0x1.0mm, FOV = 240 mm, 180 sagittal slices with interleaved acquisition). The resting-state fMRI scan used a gradient echo sequence (TR = 2000 ms, TE = 30 ms, flip angle = 75 degrees, voxel size = 4.0 x 4.0 x 4.0 mm, FOV = 256mm, 36 axial slices with interleaved acquisition).

#### 1.2 fMRI Preprocessing

The following boilerplate text is automatically automatically generated by fMRIPrep with the express intention that users should copy and paste this text into their manuscripts unchanged. It is released under the CC0 license. Results included in this manuscript come from preprocessing performed using fMRIPrep 23.2.0 (4), which is based on Nipype 1.8.6 (5).

##### 1.2.1 Anatomical data preprocessing

A total of 2 T1-weighted (T1w) images were found within the input BIDS dataset. Each T1w image was corrected for intensity non-uniformity (INU) with N4BiasFieldCorrection (Tustison et al. 2010), distributed with ANTs 2.5.0 (6). The T1w-reference was then skull-stripped with a Nipype implementation of the antsBrainExtraction.sh workflow (from ANTs), using OASIS30ANTs as target template. Brain tissue segmentation of cerebrospinal fluid (CSF), white-matter (WM) and gray-matter (GM) was performed on the brain-extracted T1w using fast (FSL (version unknown) (7). An anatomical T1w-reference map was computed after registration of 2 T1w images (after INU-correction) using mri_robust_template (FreeSurfer 7.3.2) (8). An anatomical T2w-reference map was computed after registration of 2 T2w images (after INU-correction) using mri_robust_template (FreeSurfer 7.3.2) (8). Brain surfaces were reconstructed using recon-all (FreeSurfer 7.3.2) (9), and the brain mask estimated previously was refined with a custom variation of the method to reconcile ANTs-derived and FreeSurfer-derived segmentations of the cortical gray-matter of Mindboggle (10). A T2-weighted image was used to improve pial surface refinement. Brain surfaces were reconstructed using recon-all (FreeSurfer 7.3.2) (9) and the brain mask estimated previously was refined with a custom variation of the method to reconcile ANTs-derived and FreeSurfer-derived segmentations of the cortical gray-matter of Mindboggle (10). Volume-based spatial normalization to two standard spaces (MNI152NLin6Asym, MNI152NLin2009cAsym) was performed through nonlinear registration with antsRegistration (ANTs 2.5.0), using brain-extracted versions of both T1w reference and the T1w template. The following templates were were selected for spatial normalization and accessed with TemplateFlow (23.1.0) (11): FSL's MNI ICBM 152 non-linear 6th Generation Asymmetric Average Brain Stereotaxic Registration Model [TemplateFlow ID: MNI152NLin6Asym] (12), ICBM 152 Nonlinear Asymmetrical template version 2009c p [TemplateFlow ID: MNI152NLin2009cAsym] (13).

##### 1.2.2 Preprocessing of B0 inhomogeneity mappings

A deformation field to correct for susceptibility distortions was estimated based on fMRIPrep's fieldmap-less approach. The deformation field is that resulting from co-registering the EPI reference to the same-subject T1w-reference with its intensity inverted (14,15). Registration is performed with antsRegistration (ANTs 2.5.0), and the process regularized by constraining deformation to be nonzero only along the phase-encoding direction, and modulated with an average fieldmap template (16).

##### 1.2.3 Functional data preprocessing

For each of the 1 BOLD runs found per subject (across all tasks and sessions), the following preprocessing was performed. First, a reference volume was generated, using a custom methodology of fMRIPrep, for use in head motion correction. Head-motion parameters with respect to the BOLD reference (transformation matrices, and six corresponding rotation and translation parameters) are estimated before any spatiotemporal filtering using mcflirt (FSL) (17). The estimated fieldmap was then aligned with rigid-registration to the target EPI (echo-planar imaging) reference run. The field coefficients were mapped on to the reference EPI using

the transform. The BOLD reference was then co-registered to the T1w reference using bbregister (FreeSurfer) which implements boundary-based registration (18). Co-registration was configured with six degrees of freedom. Several confounding time-series were calculated based on the preprocessed BOLD: framewise displacement (FD), DVARS and three region-wise global signals. FD was computed using two formulations following Power (absolute sum of relative motions (19) and Jenkinson (relative root mean square displacement between affines (17). FD and DVARS are calculated for each functional run, both using their implementations in Nipype (following the definitions per by Power et al. (19)). The three global signals are extracted within the CSF, the WM, and the whole-brain masks. Additionally, a set of physiological regressors were extracted to allow for component-based noise correction (CompCor) (20). Principal components are estimated after high-pass filtering the preprocessed BOLD time-series (using a discrete cosine filter with 128s cut-off) for the two CompCor variants: temporal (tCompCor) and anatomical (aCompCor). tCompCor components are then calculated from the top 2% variable voxels within the brain mask. For aCompCor, three probabilistic masks (CSF, WM and combined CSF+WM) are generated in anatomical space. The implementation differs from that of Behzadi et al. (20) in that instead of eroding the masks by 2 pixels on BOLD space, a mask of pixels that likely contain a volume fraction of GM is subtracted from the aCompCor masks. This mask is obtained by dilating a GM mask extracted from the FreeSurfer's aseg segmentation, and it ensures components are not extracted from voxels containing a minimal fraction of GM. Finally, these masks are resampled into BOLD space and binarized by thresholding at 0.99 (as in the original implementation). Components are also calculated separately within the WM and CSF masks. For each CompCor decomposition, the k components with the largest singular values are retained, such that the retained components' time series are sufficient to explain 50 percent of variance across the nuisance mask (CSF, WM, combined, or temporal). The remaining components are dropped from consideration. The head-motion estimates calculated in the correction step were also placed within the corresponding confounds file. The confound time series derived from head motion estimates and global signals were expanded with the inclusion of temporal derivatives and quadratic terms for each (21). Frames that exceeded a threshold of 0.5 mm FD or 1.5 standardized DVARS were annotated as motion outliers. Additional nuisance timeseries are calculated by means of principal components analysis of the signal found within a thin band (crown) of voxels around the edge of the brain, as proposed by (22). All resamplings can be performed with a single interpolation step by composing all the pertinent transformations (i.e. head-motion transform matrices, susceptibility distortion correction when available, and co-registrations to anatomical and output spaces). Gridded (volumetric) resamplings were performed using nitransforms, configured with cubic B-spline interpolation.
Many internal operations of fMRIPrep use Nilearn 0.10.2 (23), mostly within the functional processing workflow. For more details of the pipeline, see the section corresponding to workflows in fMRIPrep's documentation.

**1.2.4 Exclusion due to framewise displacement**
Subjects with a mean FWD of greater than 0.3mm were excluded from further analysis. Table S2 summarizes the demographic, clinical, and functional connectivity data within the subjects excluded due to high FWD (n=24; 4.5% of sample). Subjects with excess FWD were older (mean difference in age = 5.71) and had more anticorrelated target FC with the SGC seedmap (mean difference in SGC target FC = 0.08), though neither factors were not statistically significant when controlling for multiple comparisons ($p_{FDR} = 0.18$ for both).

**Table S2.** Summary of demographic, clinical, and functional connectivity data for subjects excluded due to excessive framewise displacement (n=24). *p* values in the table are uncorrected for multiple comparisons.

| | **Mean (SD) or Count (%)** | **Comparison to included subjects (chi sq or Welch's t test)** |
|---|---|---|
| Age; years | 47.83 (11.54) | $t = 2.37$ ($p = 0.03$) |
| Number of years of education; years | 16.29 (2.99) | $t = -0.97$ ($p = 0.34$) |
| Female | 13 (54%) | $\chi 2 = 0.09$ ($p = 0.92$) |
| Right-handed | 20 (83%) | $\chi 2 = 0.76$ ($p = 0.38$) |
| Baseline HRSD score | 23.08 (3.62) | $t = -0.21$ ($p = 0.84$) |
| End of rTMS HRSD score | 12.50 (7.75) | $t = -0.33$ ($p = 0.74$) |
| % change in HRSD score | -46.64 (31.45) | $t = -0.46$ ($p = 0.65$) |
| Responder | 14 (58%) | $\chi 2 = 1.13$ ($p = 0.28$) |
| Remitter | 10 (42%) | $\chi 2 = 1.78$ ($p = 0.18$) |
| Age of onset; years | 23.90 (12.6) | $t = 1.42$ ($p = 0.16$) |
| Length of current episode; months | 48.75 (65.6) | $t = 1.58$ ($p = 0.13$) |
| Anxiety comorbidity | 11 (46%) | $\chi 2 = 0.39$ ($p = 0.53$) |
| ATHF score | 7.54 (4.55) | $t = 0.71$ ($p = 0.48$) |
| 2 or more adequate trials | 12 (50%) | $\chi 2 = 0.02$ ($p = 0.88$) |
| Benzodiazepines | 5 (21%) | $\chi 2 = 0.91$ ($p = 0.34$) |
| Antidepressants | 18 (75%) | $\chi 2 = 0.11$ ($p = 0.74$) |
| Antipsychotics | 7 (29%) | $\chi 2 = 1.79$ ($p = 0.18$) |
| Theta-burst stimulation | 14 (58%) | $\chi 2 = 0.90$ ($p = 0.34$) |
| rTMS stimulation intensity | 54.96 | $t = 1.91$ ($p = 0.07$) |
| Number of daily treatments | 28.33 | $t = 0.69$ ($p = 0.50$) |
| SGC iFT Distance | 17.90 (11.57) | $t = 0.04$ ($p = 0.97$) |
| CDC iFT Distance | 17.82 (11.53) | $t = -0.45$ ($p = 0.66$) |
| SGC target FC | -0.40 (0.16) | $t = -2.34$ ($p = 0.03$) |
| CDC target FC | 0.30 (0.16) | $t = 1.94$ ($p = 0.06$) |

#### 1.3 Generating the iFTs

*Time series derivation*

For the SGC, the timeseries was derived based on a seedmap of 2,000 scans of 1,000 individuals from the Human Connectome Project (HCP; 2 scans per subject), creating a map of SGC connectivity across all gray matter voxels. Then, for each subject the SGC timeseries was computed by weighting all gray matter voxels according to the seedmap, excluding the gray matter voxels within the L-DLPFC mask. This approach has demonstrated superior signal-to-noise ratio and is a reliable reflection of individual SGC connectivity(24–26).

For the CDC, the combined CDC *r* map from Siddiqi et al. was used (27). In brief, they derived the CDC from 14 datasets of either lesions or neurostimulation target locations. For each dataset, lesion or neurostimulation target locations were used to calculate whole-brain connectivity maps. These maps were then correlated, at each voxel, with either depressive symptoms (lesions) or antidepressant effects (neurostimulation targets) to yield a 'circuit map' for each dataset, which were then combined into a single map. In the current study, to create the CDC timeseries for each subject, we weighted the signal from each gray matter voxel by the combined CDC *r* map, excluding the gray matter voxels within the L-DLPFC mask.

#### 2. RESULTS

##### 2.1 iFT Computation

The calculation of iFT distance was significantly affected by computational approach, which included the "searchlight" method or "cluster" method at 5 thresholds (SGC-iFT : $F(5,3000) = 17.29$, $p<.001$ ; CDC-iFT: $F(5,2920) = 12.33$, $p<.001$). Specifically, the mean distances for the lower cluster thresholds (0.5%, 1%, 2.5%) do not show significant post-hoc differences, however the higher cluster threshold (5%, 10%) iFTs were significantly more proximal to the group-average treatment target site for both the SGC and CDC approaches (S1A, S1C), demonstrating a decrease in inter-subject variability, similar to previous findings in scan-rescan reliability in a non-clinical sample (24). Finally, the searchlight method returned significantly greater distances from the treatment target, compared to all cluster thresholds, for the SGC-iFT approach (S1A, S1C). Spearman rho values for correlation with changes in HRSD scores were non-significant at all threshold levels (S1B, S1D).

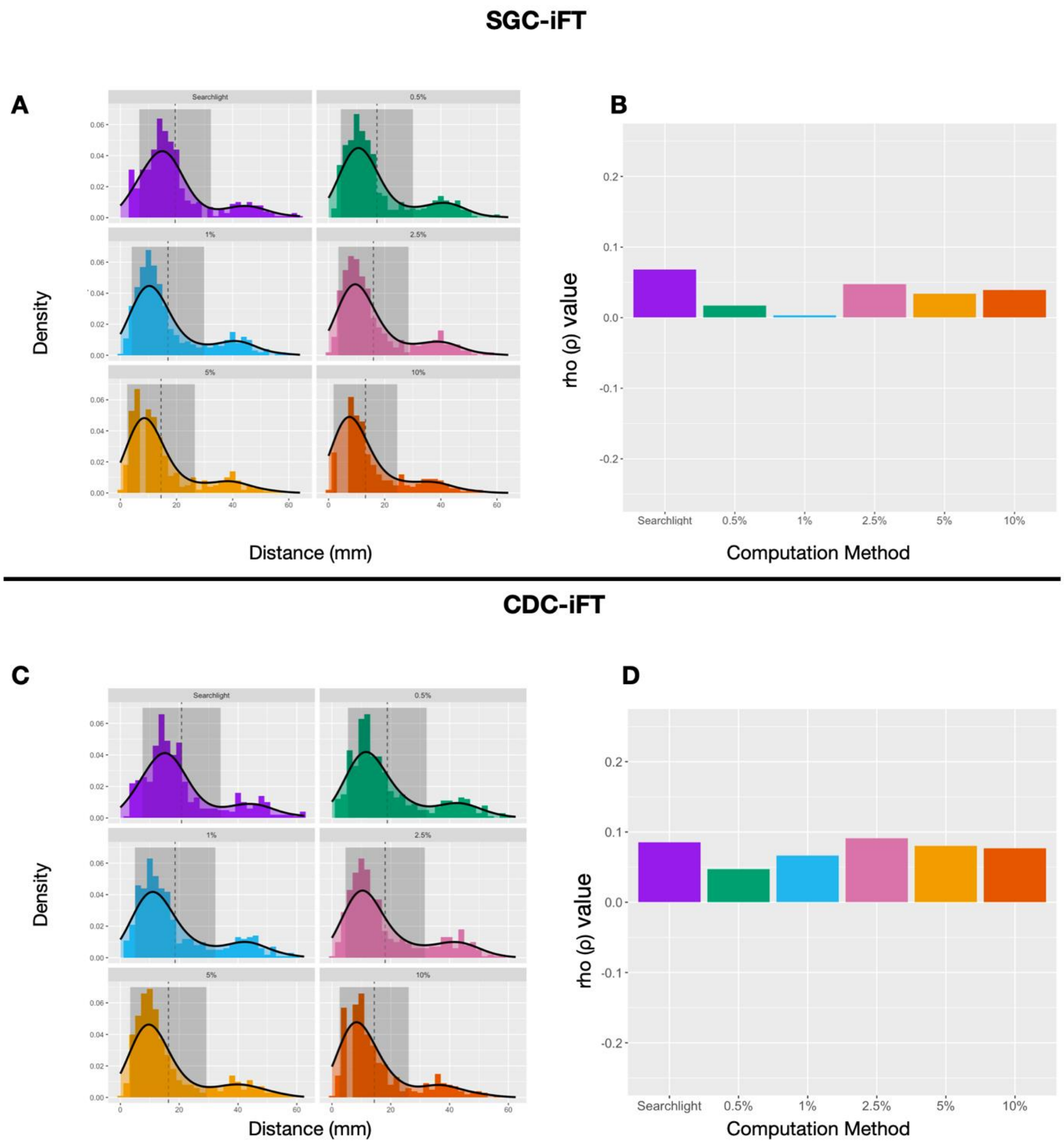

**Figure S1.** Effects of computation methods on iFT results. (A) Density SGC-iFT distances to the treatment target using the searchlight approach and varying cluster thresholds. Vertical dashed lines represent the mean distance, while shaded grey represents standard deviation.(B) Correlations (rho (ρ) values) between SGC-iFT distance and symptom improvement do not change significantly across computation method, all of which fell between 0.003 – 0.07. (C) Density CDC-iFT distances to the treatment target across cluster thresholds. Vertical dashed lines represent the mean distance, while shaded grey represents standard deviation. (D) Correlations (rho (ρ) values) between CDC-iFT distance and symptom improvement do not change significantly with varying cluster thresholds, all of which fell between 0.05 – 0.09.

#### 2.2 Effect of data quality on results

tSNR of the fMRI signal was calculated after minimal preprocessing in fMRIPrep, but prior to ICA-AROMA, nuisance signal regression, and bandpass filtering. tSNR was calculated across all grey matter voxels (after dropping the first 4 volumes to achieve steady-state) using the following equation:

$$tSNR = \frac{mean(GreyMatter)}{SD(GreyMatter)}$$

The mean tSNR of the sample was 87.87 (SD=18.63, range = 38.10-132.57). There was no relationship between tSNR and distance values (SGC iFT: $\rho = -0.05$, $p = 0.28$; CDC iFT: $\rho = -0.04$, $p = 0.40$). The relationship between distance and treatment outcomes was not altered when controlling for tSNR, remaining non-significant for both iFTs (SGC-iFT: $\rho' = 0.02$, $p = 0.61$; CDC-iFT: $\rho' = 0.05$, $p = 0.28$).

To assess whether null findings may be driven by low tSNR, we assessed the differences in the relationship of both distance and FC to treatment outcomes across tSNR quartiles ($1^{st}$ quartile:38.10 - 74.57; $2^{nd}$ quartile: 74.66 – 88.16; $3^{rd}$ quartile: 88.18 – 101.18; $4^{th}$ quartile: 101.29 – 132.57). Within all quartiles, there was no significant relationship between distance and treatment outcomes or between target FC and treatment outcomes, as shown in Figure S2.

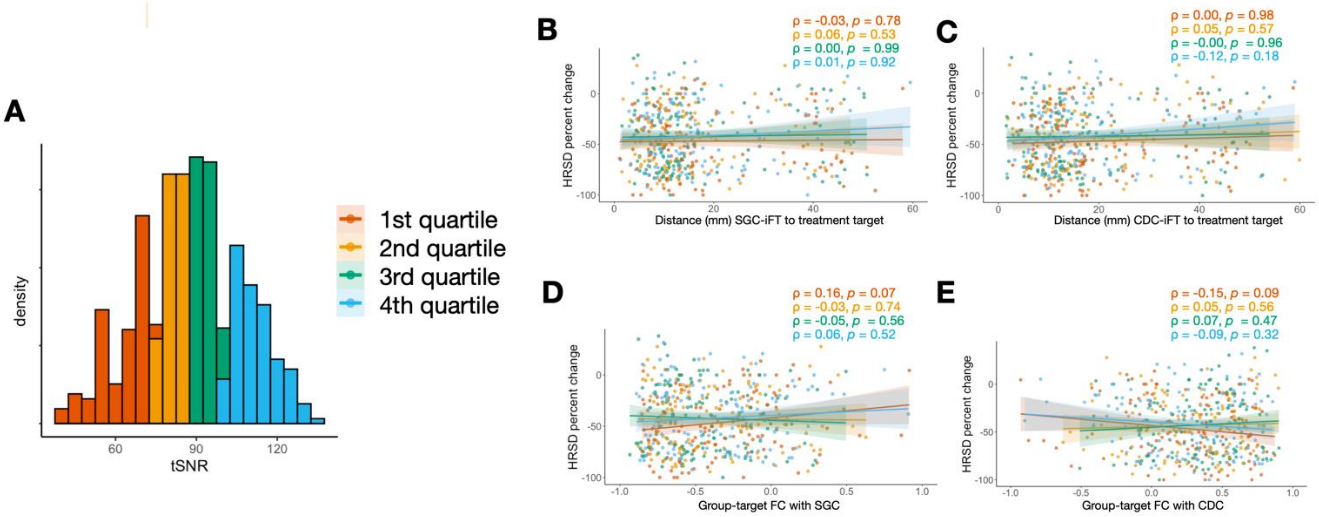

**Figure S2.** (A) Subjects were split into quartiles based on tSNR to assess whether associations between distance and/or target FC varied as a function of data quality. There was no relationship between iFT distance to treatment target and clinical response across any of the tSNR quartiles for both the SGC-iFT (B) and the CDC-iFT (C). With respect to target FC, there was also no relationship between D) target anticorrelation with the SGC-iFT and E) target correlation with the CDC and clinical response across all quartiles. **ρ** (rho), ***p*** (p value)

We also assessed tSNR within the SGC, given the challenges related to signal dropout in this region (28,29). The seedmap approach for generating SGC timeseries was developed to migitate these challenges(28), and reliably reflect ground-truth SGC signal, as inferred from scanning sequences that are able to capture high-quality signal from this region(25).

After the fMRIPrep pipeline, but prior to nuisance regression, smoothing, and bandpass filtering, the mean tSNR within the SGC ROI (10mm radius sphere at [6, 16, -10]) was 48.20 (SD = 23.41). A brain-wide map of tSNR across voxels is shown in Figure S3 (A), along with the distribution of SGC ROI tSNR within the sample (B), and examples of single-subject tSNR plots showcasing a subject with low SGC tSNR (C) and high SGC tSNR (D).

To determine any effects of SGC signal dropout on our findings, we performed sensitivity analyses, looking at whether effects were detectable in subjects with a SGC ROI tSNR of 50 or greater (n = 211, 42% of sample) or in subjects with a SGC ROI tSNR of 80 or greater (n = 52, 10% of sample).

Subjects with an SGC ROI tSNR of 50 or more showed no effect of iFT proximity (SGC $\rho = -0.01$, $p = 0.88$; CDC $\rho = 0.07$, $p = 0.33$) or target FC (SGC $\rho = 0.07$, $p = 0.34$; CDC $\rho = -0.01$, $p = 0.86$) on treatment outcomes. Similar findings were seen in subjects with an SGC ROI tSNR of 80 or more, with no effect of iFT proximity (SGC $\rho = -0.02$, $p = 0.89$; CDC $\rho = -0.01$, $p = 0.96$) or target FC (SGC $\rho = -0.08$, $p = 0.58$; CDC $\rho = 0.13$, $p = 0.35$) on treatment outcomes.

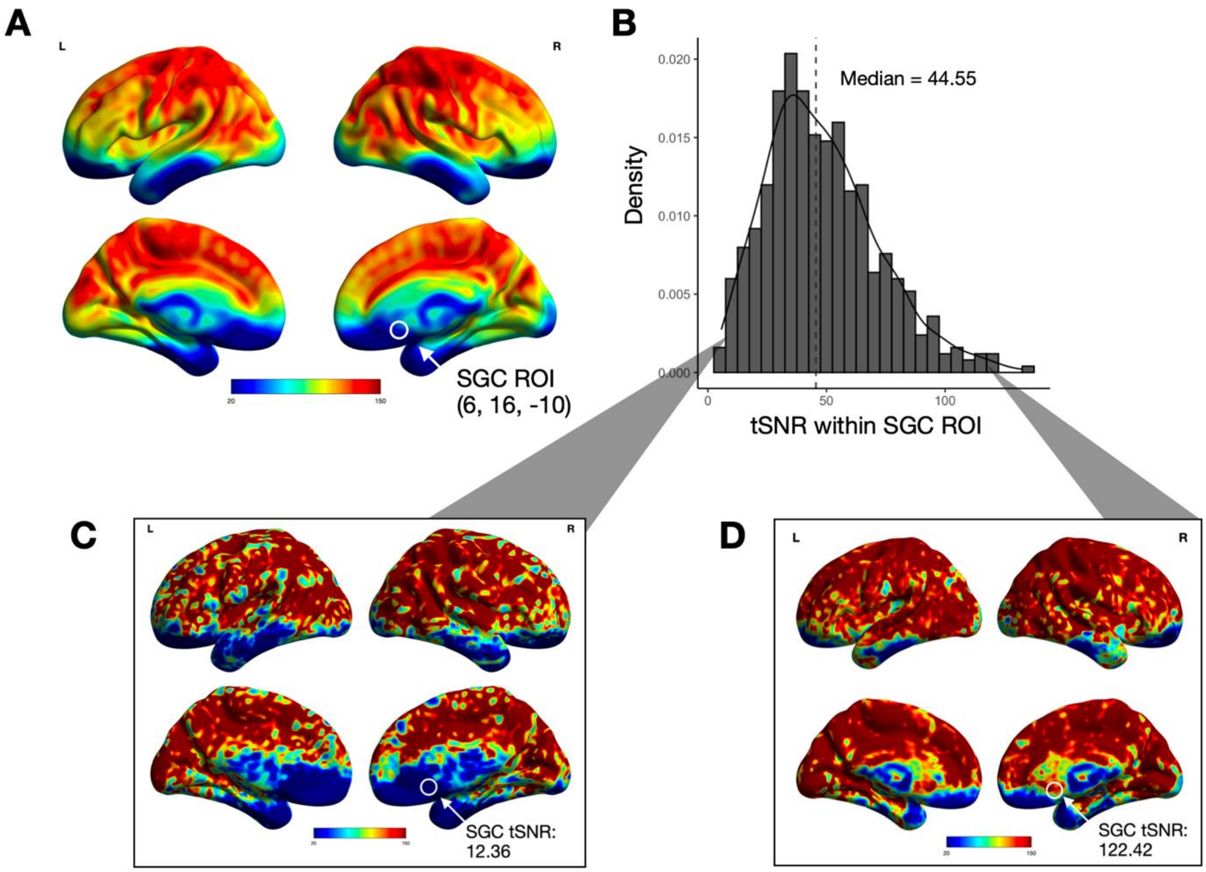

**Figure S3.** (A) Mean tSNR across the brain in the entire sample. Signal dropout was highest in orbitofrontal and inferior temporal regions. The SGC ROI is circled for clarity (M = 48.20, SD =

23.41 in this region) (B) Distribution of tSNR values within the SGC ROI. (C) Example of tSNR map in a subject with poor signal within the SGC ROI. (D) Example of tSNR map in a subject with good signal within the SGC ROI.

#### 2.3 Effect of Scanner Site & Scan Protocol

Since CAMH was only used in the CARTBIND cohort, there were 5 groups for comparison. For both SGC-iFT and CDC-iFT, there was a significant effect of site/scan protocol.

- SGC-iFT: $F(4,496) = 9.01$, $p < 0.001$. Subjects scanned with the CARTBIND protocols at the UHN and CAMH sites had significantly higher distances (M=23.87, SD=15.15 and M=20.68, SD=13.57) compared to subjects scanned at UBC (M=13.88, SD=11.89); $p = 0.04$ and $p = 0.003$. For the THREED protocols, we found that UHN distances were greater (M=16.88, SD=12.46) compared to the UBC site (M=10.41, SD=11.89), $p = 0.01$. For the sites which had scanned both cohorts, we found that the CARTBIND protocol was associated with significantly greater distances compared to the THREED protocol for the UHN site only (UHN: $p = 0.01$; UBC: $p = 0.69$).
- CDC-iFT: $F(4, 496) = 10.45$, $p < 0.001$. Subjects scanned with the CARTBIND protocols at the UHN and CAMH sites had significantly higher distances (M=26.25, SD=14.81 and M=23.64, SD=14.73) compared to subjects scanned at UBC (M=13.12, SD=10.51); $p < 0.001$ and $p = 0.003$. For the THREED protocols, we found no difference in distance across sites (UHN: M=17.97, SD=12.76, UBC: M=13.92, SD=9.92), $p = 0.26$. For the sites which had scanned both cohorts, we found that the CARTBIND protocol was associated with significantly greater distances compared to the THREED protocol for the UHN site only (UHN: $p = 0.001$; UBC: $p = 0.99$).

Multiple linear regressions, accounting for the interaction between site/scan protocol and distance, found no significant effect of iFT distance on HRSD scores for the SGC-iFT (distance: $F(1,491) = 1.39$, $p = 0.24$; distance x site/scan protocol: $F(4,491) = 1.13$, $p = 0.34$) or CDC-iFT (distance: F(1,491 = 2.48, $p = 0.12$; distance x site/scan protocol: $F(4, 491) = 0.21$, $p = 0.93$). The relationships between distance and HRSD change, stratified by site/scan protocol, are shown in Figures S4C and S4D.

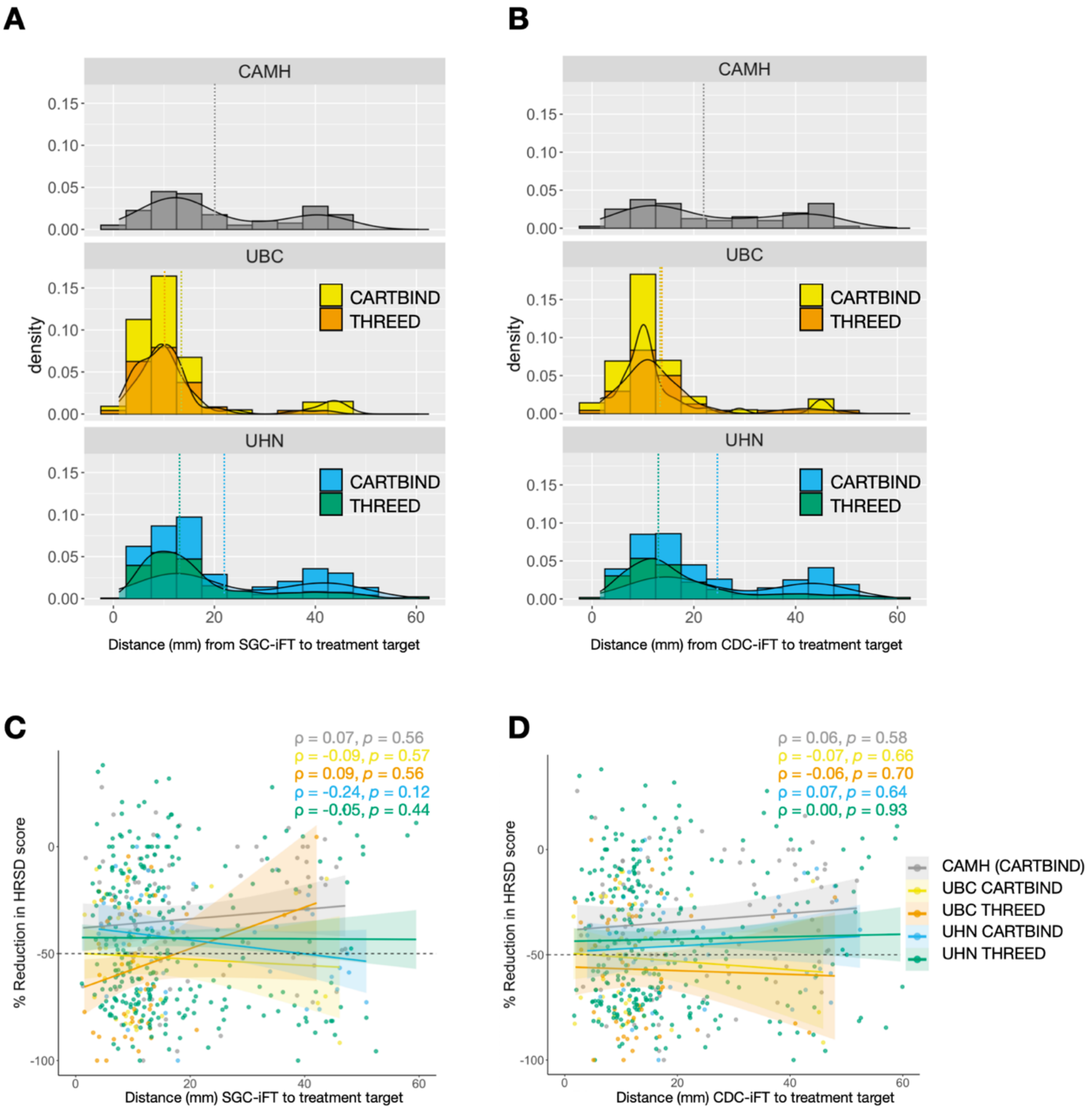

**Figure S4.** Differences in scanner site and scan protocol did not have a significant influence on findings. (A) Shows the distribution of distances calculated for the SGC-iFT across sites, and protocol for sites that scanned both cohorts. (B) Shows the distribution of distances calculated for the CDC-iFT. For both (A) and (B), site means are plotted as horizontal dashed lines. (C) Scatterplots of the relationship between SGC-iFT distance and treatment response stratified by site/scan protocol. All correlations were non-significant (ρ ranging from -0.24 to 0.09, *p* (uncorrected) ranging from 0.12 – 0.57). (D) Scatterplots of the relationship between CDC-iFT distance and treatment response stratified by site. All correlations were non-significant. **ρ** (rho), ***p*** (p value)

#### 2.4 Effect of Cohort and Treatment Allocation

We assessed whether there were different effects of iFT proximity on treatment outcomes within the 2 cohorts included in the analysis. Correlations between proximity and outcomes remain

small and non-significant for both the THREE-D cohort (SGC-iFT $\rho = 0.01$ , $p = 0.96$; CDC-iFT $\rho = 0.01$, $p = 0.82$) and the CARTBIND cohort (SGC-iFT $\rho = 0.01$, $p = 0.95$; CDC-iFT $\rho = 0.09$, $p = 0.26$).
When looking at effects across treatment allocations within each clinical trial (10Hz or iTBS for THREE-D; once or twice daily treatments for CARTBIND), there was a weak correlation between CDC-iFT proximity and treatment outcomes for the twice-daily CARTBIND group only, which did not survive correction for multiple comparisons ($p_{FDR} = 0.32$).
Treatment allocation effects are summarized in Figure S5.

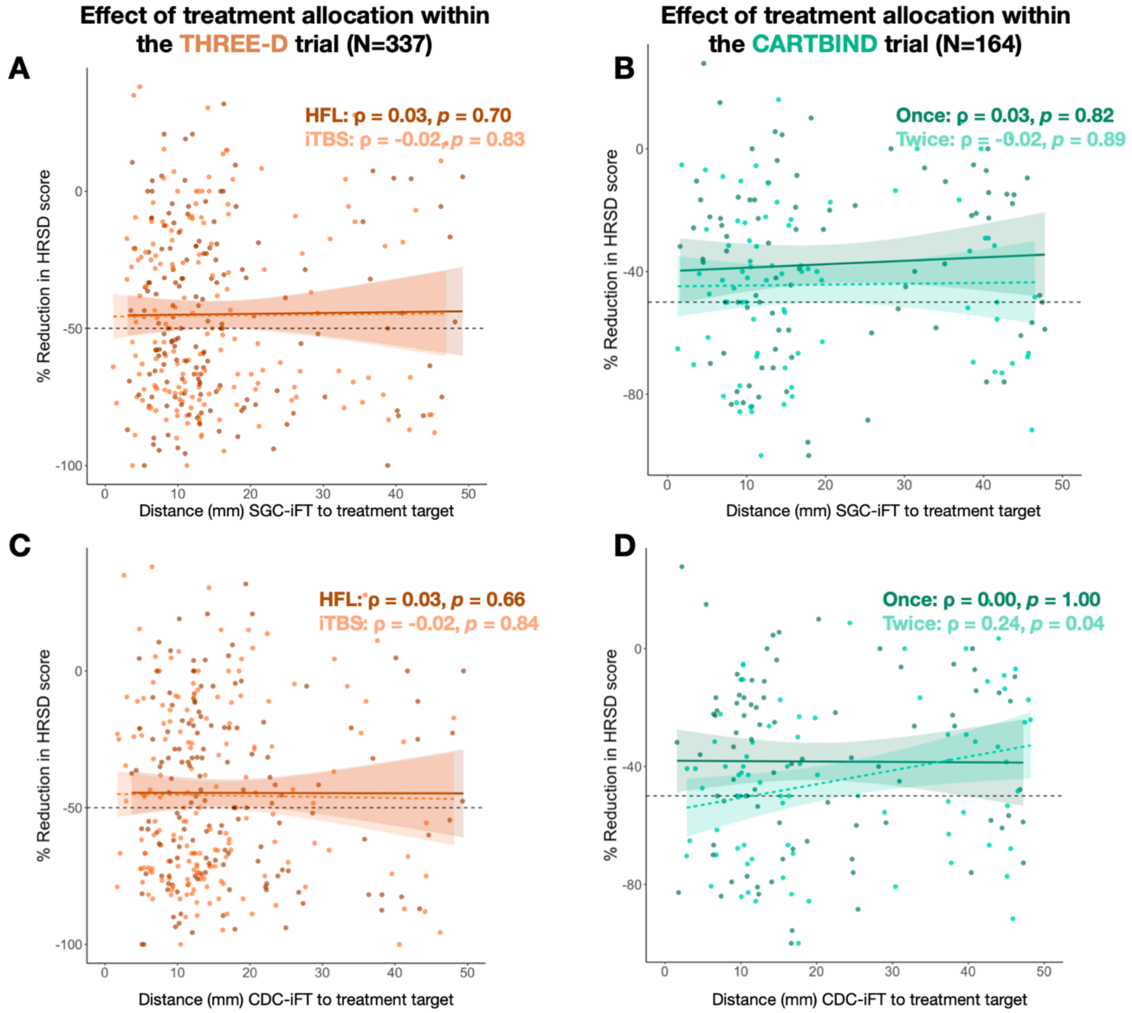

**Figure S5.** The relationship between proximity to iFTs and treatment outcomes was the same across cohorts (*ie* clinical trials) and treatment allocations (*ie* what treatment each participant received within their respective trial). This is demonstrated for THREE-D in (A) for SGC-iFT and (B) for CDC-iFT. Similarly, CARTBIND is demonstrated in (C) for SGC-iFT and (D) for CDC-iFT. *p* values represented in the figure are uncorrected for multiple comparisons. ρ (rho), ***p*** (p value)

#### 2.6 Changes in HRSD-17 symptom subdomains

We conducted post-hoc exploration of the relationship between iFT proximity and treatment outcomes relating to change in symptom cluster scores (follow-up minus baseline) and iFT target proximity. Cluster scores were computed as the sum of all HRSD items loading into the factor multiplied by their respective factor loadings (30). Difference scores, rather than % change, were computed, as instances where symptoms worsened resulted in very high percent changes. Instead, partial correlations were computed with baseline scores as the z variable to control for the effect of baseline score on change.

No correlations were significant, as summarized in Figure S6.

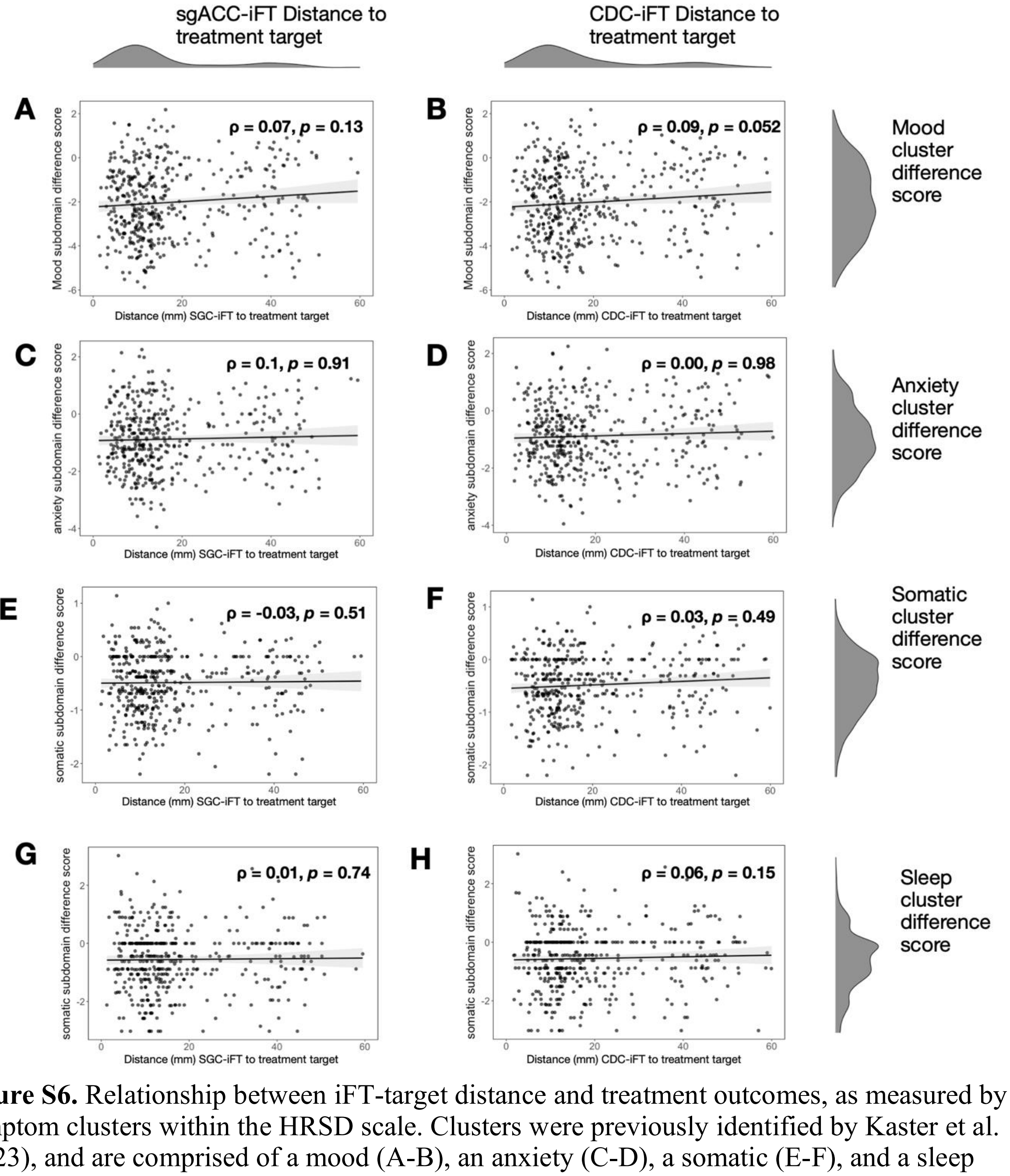

**Figure S6.** Relationship between iFT-target distance and treatment outcomes, as measured by symptom clusters within the HRSD scale. Clusters were previously identified by Kaster et al. (2023), and are comprised of a mood (A-B), an anxiety (C-D), a somatic (E-F), and a sleep cluster (G-H). No correlations were significant. *p* values are uncorrected for multiple comparisons. ρ (rho), *p* (p value).

#### 2.7 Counterfactual analyses of iFT distance to Beam F3 and "5cm rule" and functional connectivity

To explore the counterfactual scenario where heuristic targeting was used, we computed the distance between both iFTs and either of the heuristic targets (Beam F3 or 5cm rule). We also computed FC with the SGC and CDC timeseries at each of these targets, using the 12mm weighted cone approach.

For these heuristic targets, we relied on previously reported average coordinates using MNI coordinates [-41, 16, 54] for the 5cm target and [-43, 46, 32] for the BeamF3 target (31). In Euclidean distance, the MNI coordinates for the group-based target [-38, 44, 26] was 39mm from the average 5cm target and 8mm away from the average BeamF3 target.

Visualization of the three targets (stimulation target, average 5cm, and average BeamF3) are shown in Figure S7.

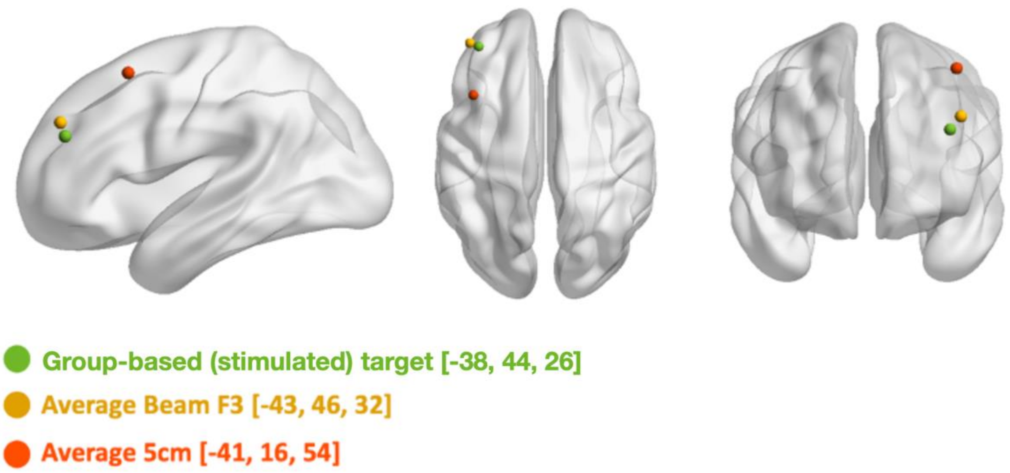

**Figure S7.** Group-based (stimulated) target, average BeamF3, and average 5cm targets projected onto cortical surface.

Table S3 summarizes iFT distances and FC with the SGC or CDC for each target, as well as results of repeated measures ANOVAs (performed with *r* package "rstatix") comparing these values across the three targets, with post-hoc pairwise comparison performed with Bonferroni correction for multiple comparisons.

Firstly, iFTs were significantly further from the heuristic-based targets compared to the group-based target. For SGC-iFTs, pairwise comparisons revealed that the average 5cm target was associated with significantly greater distances compared to the averageBeamF3 target (t(500)=10.3, *p*<.001) and the group-based target (t(500)= 14.0, *p*<.001), while the average BeamF3 target was significantly further than the group-based target (t(500)=6.53, *p*<.001). For CDC-iFTs, the same pattern emerged, with the average 5cm target again being the furthest compared to both the average BeamF3 target (t(500)=10.6, *p*<.001) and the group-based target (t(500)=11.67, *p*<.001) and the averageBeamF3 target being further than the group-based target (t(500)=16.9, *p*<.001).

In line with findings of the group-based target being more proximal to iFTs, computed FC at each target showed a similar pattern. For FC with the SGC, post-hoc pairwise corrections revealed that the average 5cm target showed the weakest values compared to the average BeamF3 target (t(500)=11.9, $p$<.001) and the group-based target (t(500)=17.6, $p$<.001), while BeamF3 was also associated with weaker FC (t(500)=16.0, $p$<.001) compared to the group-based target. For FC with the CDC, similarly the average 5cm target had weaker correlations compared to the average BeamF3 target (t(500)=-15.7, $p$<.001) and the group-based target (t(500)=-17.6, $p$<.001), again with the BeamF3 target also showing reduced FC compared to the group-based target (t(500)=-11.3, $p$<.001).

**Table S3.** Summary of mean values of iFT distance and functional connectivity (within a 12mm weighted cone) at each of the selected targets. All F statistics were significant at $p$<.001. All targets were statistically different (post-hoc pairwise comparisons) in terms of distance and FC values at $p$<.001.

| | **Average 5cm target** | **Average BeamF3 target** | **Group-based target** | **F test (DFn, DFd)** |
|---|---|---|---|---|
| | **Mean (SD)** | **Mean (SD)** | **Mean (SD)** | |
| SGC-iFT distance | 30.85 (12.88) | 20.61 (11.69) | 17.24 (12.87) | 137.22 (1.42, 710.27) |
| CDC-iFT distance | 31.43 (12.91) | 21.00 (11.87) | 18.83 (13.37) | 126.59 (1.01,506.12) |
| SGC FC | -0.07 (0.18) | -0.21 (0.18) | -0.39 (0.24) | 251.47 (1.38, 692.26) |
| CDC FC | 0.00 (0.18) | 0.15 (0.17) | 0.29 (0.23) | 260.46 (1.38, 690.55) |
